# Eye Blinks as a Visual Processing Stage

**DOI:** 10.1101/2023.06.18.545489

**Authors:** Bin Yang, Janis Intoy, Michele Rucci

## Abstract

Humans blink their eyes frequently during normal viewing, more often than it seems necessary for keeping the cornea well lubricated. Since the closure of the eyelid disrupts the image on the retina, eye blinks are commonly assumed to be detrimental to visual processing. However, blinks also provide luminance modulations rich in spatial information to neural pathways highly sensitive to temporal changes. Here we report that the luminance transients from blinks enhance visual sensitivity. By coupling high-resolution eye-tracking in human observers with modeling of blink transients and spectral analysis of visual input signals, we show that blinking increases the power of retinal stimulation and that this effect significantly enhances visibility despite the time lost in exposure to the external scene. We further show that, as predicted from the spectral content of input signals, this enhancement is selective for stimuli at low spatial frequencies and occurs irrespective of whether the luminance transients are actively generated or passively experienced. These findings indicate that, like eye movements, blinking acts as a computational tool in a visual processing strategy that uses motor behavior to reformat spatial information in the temporal domain.

## Introduction

Humans blink their eyes every few seconds. While the frequency of blinks varies greatly across individuals and tasks^1–3^, blinking is critically important: it maintains the eye well lubricated^4, 5^, regulating the precorneal tear film^6^ and improving the optical quality of the image on the retina^7^. In exerting these beneficial actions, however, blinks challenge visual processing, as they temporarily occlude the external scene. Since each blink can last as long as 300 ms^8–10^, blinking significantly disrupts the acquisition of visual information and may substantially delay motor responses to important visual events.

This problem is further exacerbated by the attenuation in visual sensitivity that occurs around the time of an eye blink^8, 9, 11^. Humans are normally not aware of the interruptions imposed by blinks in the visual stream entering the eye. This is a remarkable accomplishment considering that comparable input changes would be startlingly obvious if resulting from the external scene rather than eye blinks. This process seems to be partly mediated by an internal suppression mechanism that attenuates visual sensitivity, akin to the one occurring during saccadic eye movements^12–18^. Since this suppression precedes and outlasts each blink^13^, it leads to reduced visibility lasting even longer than the blink duration.

These considerations, together with the observation that humans blink more often than necessary for keeping the eye well lubricated^6, 7, 19^, suggest that eye blinks must also serve other functions. In principle, there are two complementary ways in which blinks could contribute to visual perception: by intervening in visual processing via their associated extraretinal signals; and by directly affecting neural responses via the luminance changes they cause on the retina. Whereas the former possibility has been the subject of recent investigations^20, 21^, the perceptual consequences of the luminance modulations exerted by blinks have been only marginally investigated.

Yet, several considerations suggest that these signals could be beneficial. Given that neurons in the early visual system tend to be strongly sensitive to input changes^22–24^, one would expect blinks to sharply modulate neural activity. Indeed, transient responses have been observed in the visual cortex at the time of eye blinks, with activity first decreasing as the eyelids close and then rapidly recovering at the reappearance of the stimulus^25, 26^. Notably, activity rebounds to a higher level immediately following a blink^26^, an effect driven by the re-afferent stimulation that may facilitate visual encoding.

Although with different characteristics, the luminance modulations caused by other types of motor actions, eye movements, play critical roles in visual perception^27–31^. Both saccades^32–35^ and ocular drifts^36, 37^—the incessant fixational motion of the eye in between saccades^38–40^— yield luminance transients that are perceptually beneficial. These signals enhance vision in complementary ways, as expected from their specific characteristics, *i.e.*, the distinct ways saccades and drifts transform an external spatial scene into a spatiotemporal flow onto the retina. The abrupt luminance changes delivered by blinks differ considerably from the modulations induced by eye movements. But, as it happens for saccades^41^, one may expect that, in a sufficiently low spatial frequency range, blinks will yield a more effective input than that normally present during fixation.

Here, we focus on the perceptual consequences of the luminance changes resulting from blinks. We show that these modulations enhance contrast sensitivity as predicted by their spectral characteristics, specifically the power within the temporal range of retinal sensitivity. We further show that this enhancement also occurs during passive exposure to similar transients, even if they are not actively generated by blinks.

## Methods

### Subjects

Data were collected from 12 subjects (4 females and 8 males; average age 22 years). Nine and 6 subjects participated, respectively, in the experiments of Figs. 2 and 5, and 2 subjects took part in the control experiment of Fig. 4. All subjects were paid to participate and possessed normal emmetropic vision, as assessed via a standard eye-chart test. With the exception of one of the authors, all subjects were näıve about the purposes of the experiments. Informed consent was obtained from every subject following the procedures approved by the Research Subjects Review Board at the University of Rochester.

### Stimuli and Apparatus

Stimuli consisted of gratings displayed over a visual field of 21.2^◦^ × 11.9^◦^ for 2.5 s. The frequency of the gratings varied across experiments. It was 3 cycles/degree (cpd) in the experiments of Fig. 2 and Fig. 5; and 1 or 10 cpd (randomly interleaved across trails) in the experiment of Fig. 4. In all experiments, the gratings varied randomly across trials in both orientation (±45° relative to the vertical meridian) and phase (0°, 90°, 180°, or 270°). Stimuli were displayed on a calibrated LCD monitor (Acer Predator XB272) at a 200 Hz refresh rate and at a resolution of 1366 × 768 pixels. They were observed binocularly from a distance of about 160 cm, with each pixel subtending *∼*1^t^. The head of the observer was immobilized by means of a head-rest and a custom dental-imprint bite bar.

The movement of the right eye was continually monitored by a Dual Purkinje Image eye-tracker, either the analog commercial device (a generation 6 DPI; Fourward Technology) or an in-house developed digital apparatus (the dDPI^42^). Both systems operate by measuring the relative displacement between the first and fourth Purkinje reflections of an infrared beam and provide sub-arcminute resolution with artificial eyes. In the DPI, the analog oculomotor signal was first low-pass filtered at 500 Hz and then sampled at 1 kHz. The dDPI directly delivers digital measurements at either 331 Hz or 1 kHz. We measured only one eye because of the conjugacy of eye blinks^43^.

### Experimental Procedures

Data were collected in blocks of 50 trials. A typical experimental session consisted of 5 blocks, lasting approximately 1 hour. Before each block of trials, the eye-tracker was tuned for optimal tracking and calibrated following standard procedures described in previous publications^44, 45^. Breaks in between blocks allowed the subject to rest.

Subjects were asked to report whether a grating was tilted 45° clockwise or counter-clockwise. Each trial started with the subject maintaining fixation at the center of the monitor, indicated by four arches (radius 0.5°) on a uniformly dark gray field (luminance 3.8 cd*/*m^2^; Fig. 2A). After a random interval of 900-1100 ms, the contrast of the grating gradually increased reaching an individually predetermined level over a period of 1.5 s. It then remained constant for an additional 1 s. A high-contrast white-noise mask was then displayed for 1 s over the entire monitor to end the trial, and the observer entered their perceptual response by pressing one of two buttons on a joypad.

In each trial, an auditory cue (a 50-ms beep) instructed the subject to execute a blink. In half of the trials the cue occurred during the presentation of the stimulus, 600 ms after the onset of the contrast ramp (the Stimulus-Blink condition). In the other half of the trials, the cue was given 800 ms before the ramp onset, so that the blink occurred when the stimulus was not present (No-Stimulus-Blink condition). Trials from the two conditions were randomly interleaved within each block.

The specific contrast value reached by the stimulus was adjusted for each individual subject to obtain *∼*80% correct responses. This was achieved in a preliminary calibration procedure via the PEST method^46^. In some cases, due to variability in the subject’s performance, the contrast was slightly adjusted across experimental sessions to remain close to threshold. These small adjustments always occurred in both Stimulus- and No-Stimulus-Blink conditions, so that the contrast was maintained identical in the two sets of trials.

In the experiment of Fig. 5, rather than executing a blink, subjects were exposed to changes in the stimulus that simulated the luminance transients caused by blinks. Specifically, the luminance of the monitor was transiently minimized (0.02 cd*/*m^2^) following a time course similar to that of an eye blink. Both the time of occurrence of a simulated blink and its duration were randomly sampled from their corresponding distributions in the blink data collected in the experiments.

### Data Analysis

Recorded eye traces were segmented into complementary periods of blinks, saccades, and drifts. Blinks were reliably detected from the disappearance of the first Purkinje reflection of the infrared beam delivered by the eye-tracker, which in our apparatus only happens when the cornea is covered by the palpebrae. Periods in which the eye moved faster than 3°*/s* were labeled as saccades or microsaccades based on the displacement amplitude, whether greater or smaller than 30^t^, respectively. Only trials with optimal eye-tracking and in which subjects did not execute non-prompted blinks were selected for data analysis. Furthermore, to avoid other sources of luminance modulations, in the experiments of Figs. 2 and 5, we discarded all trials that contained saccades of any amplitude, including microsaccades. Thus, in the trials selected for data analysis, the eye only moved by means of ocular drift and contained only one blink (Fig. 2) or no blinks at all (Fig. 5). The number of selected trials in both experiments ranged from 141 to 563 across subjects.

To investigate the visual consequences of blinks, we compared performance between the Stimulus-Blink trials and the No-Stimulus-Blink trials. Performance was quantified by means of both proportions of correct responses and discriminability index, *d*^t^. Since *d*^t^ goes to infinity when the hit or false alarm rates get close to 0 or 1, we truncated the hit and false alarm rate between 0.01 and 0.99, corresponding to a maximum *d*^t^ of 4.65^47^. Differences in performance were evaluated both on average across subjects and individually for each subject. The probability density distributions of blink reaction times (the delay between the onset of the auditory cue and the onset of blink; Fig. 2B) and blink durations (Fig. 2C) were estimated over 50-ms bins.

### Spectral Estimation

To examine the spatio-temporal signals resulting from eye blinks, we reconstructed the luminance flow impinging onto the retina and estimated its power within the range of human temporal sensitivity. Let *I*(***x***) denote the contrast of the stimulus as a function of space ***x***. The visual flow *L_I,_****_ξ_***(***x****, t*) resulting from observing *I*(***x***) during a sequence of eye movements ***ξ***(*t*) and blinks can be expressed as:

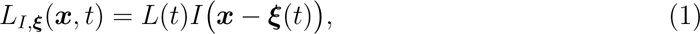

where *t* indicates time and *L*(*t*) models the global temporal modulations resulting from blinks and other factors. In our case, *L*(*t*) = *A*(*t*)*B*(*t*), where *A*(*t*) represents the time-varying contrast profile of the stimulus on the display, the 1.5 s ramp followed by the 1 s plateau. The term *B*(*t*) is a binary function that models the occurrence of an eye blink (0 during the blink, 1 otherwise).

The power spectrum of the visual input was estimated by means of the periodogram:

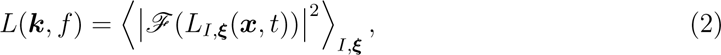

where *ℱ* represents the Fourier Transform operator, 〈〉_𝛪,*ξ*_ indicates averaging across stimuli and recorded eye trajectories, and ***k*** and *f* are the spatial and temporal frequencies. We used the spatio-temporal factorization approach proposed in previous studies^41, 48^. This method enhances spectral resolution by assuming that the image on the retina and its motion are independent—a plausible assumption on average across the visual field:

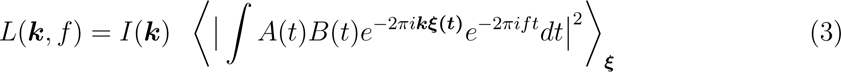

where *I*(***k***) is the power spectrum of the stimulus and the second term represents the temporal redistribution of power caused by eye movements, blinks, and the dynamics of stimulus presentation. In our experiments, we used a grating at spatial frequency *k*_0_: *I*(***x***) = sin (2*πk*_0_***x***).

To understand how this input signal may affect perception, we computed its power within the temporal range of human sensitivity. To this end, *L*(***k****, f*) was filtered by the known temporal sensitivity function of the human visual system^49^, *H*(*f*). That is, we first weighted *L*(***k****, f*) by *H*(*f*) and then integrated across temporal frequencies:

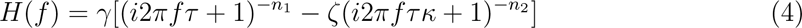

where *κ* = 1.33, *n*_1_ = 9, *n*_2_ = 10, *τ* = 4.94, *ζ* = 1, and *γ* = 20.

In Fig. 1, to examine how blinks alter the luminance flow impinging onto the retina during normal fixational instability, we compared the power of visual input signals delivered by gratings at various spatial frequencies in the presence and absence of blinks. Ocular drift (the term ***ξ***(*t*) in Eq. 1) was modeled as Brownian motion^48, 50^ with a diffusion constant of 10 arcmin^2^/s^36, 37^. Blink duration (the term *B*(*t*)) followed a normal distribution with a mean of 150 ms and a standard deviation of 10 ms.

**Figure 1:**
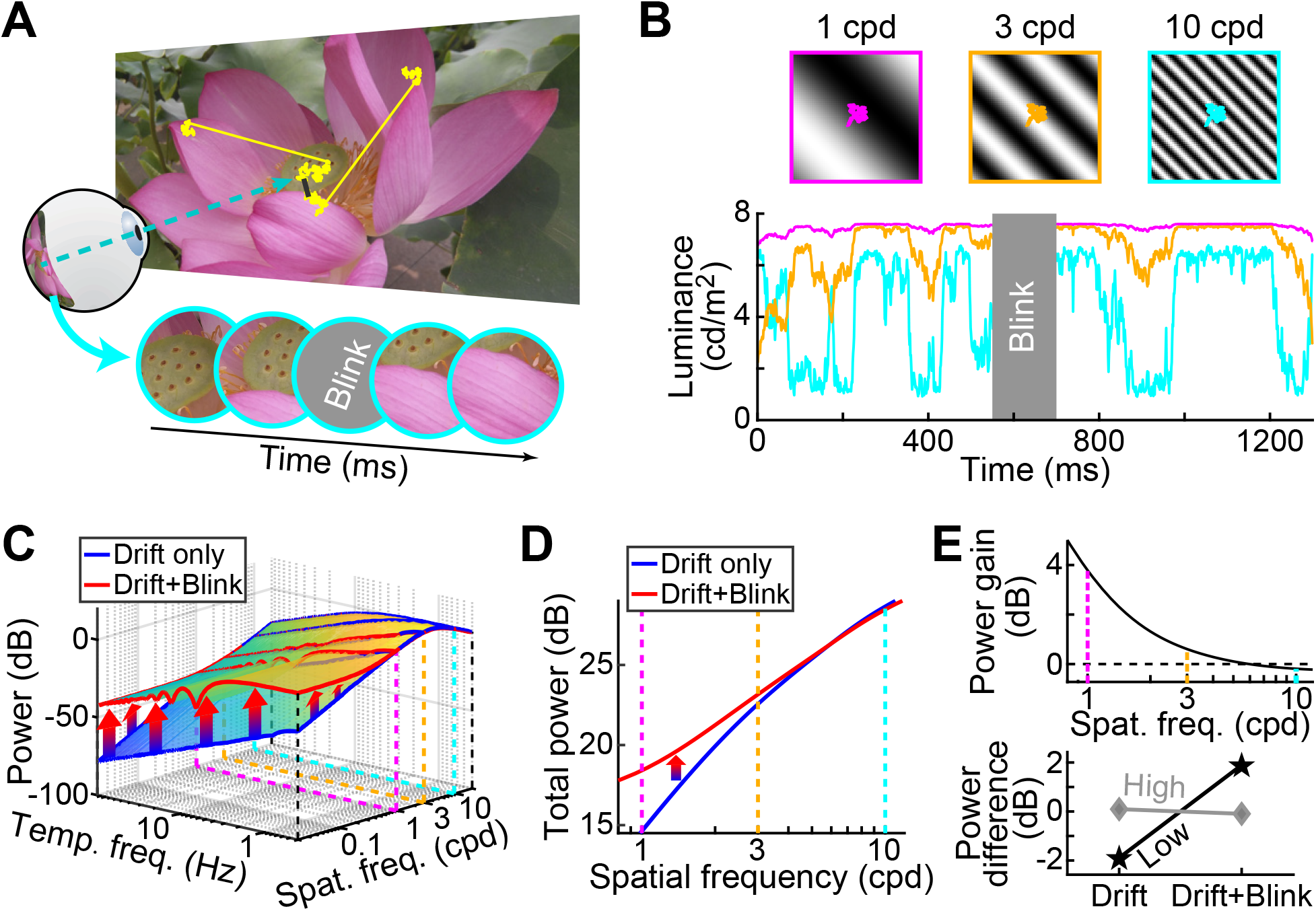
Predicted consequences of blink transients. (***A***) A sequence of eye movements (yellow trace) is interrupted by a blink (dark segment), which transiently occludes the visual input to the retina (Bottom). (***B***) Luminance modulations experienced by a small retinal area (3^t^ diameter) during exposure to the same eye drift trajectory over gratings at three different spatial frequencies. Input signals become larger and faster with increasing spatial frequency. (***C***) Power spectra of simulated visual input in the presence (Drift+Blink; red lines) and absence of eye blinks (Drift only; blue). Ocular drift was modeled as Brownian motion and blinks as transient periods of zero contrast. (***D***) Power of the resulting input signals within the temporal range of human sensitivity. (***E***) Blinks are expected to increase the power up to *∼*5 cpd (Top panel), yielding stronger visual input signals at low (Bottom; black line, 1 cpd) but not high spatial frequencies (gray, 10 cpd).

## Results

To predict the visual consequences of eye blinks, we first examined how their occurrence alters the luminance flow normally impinging onto the retina (Fig. 1A). In the fixation periods in between saccades, ocular drifts continually modulate visual input signals. These modulations depend on the stimulus, increasing both in amplitude and speed as the spatial frequency of the stimulus increases^37, 51^ (Fig. 1B). In contrast, the modulations resulting from blinks do not depend on spatial frequency, as blinks deliver signals with identical dynamics irrespective of the stimulus. Thus, if blink transients are used by the visual system, one may expect them to exert a stronger influence in the low spatial frequency range, where the modulations resulting from fixational eye movements are smaller.

We quantitatively explored this idea in simulations of the visual input signals present during fixation (Fig. 1C-E). We modeled the fixational eye movements normally performed by healthy observers as Brownian motion^48, 50^, while blinks acted by transiently suppressing the contrast of the resulting spatiotemporal luminance flow. Fig. 1C shows the power spectrum of the simulated visual input signal during observation of a stationary white noise stimulus, *i.e.*, a stimulus that contains all spatial frequencies with equal amplitude. As shown by these data, the occurrence of an eye blink greatly increases power at low spatial frequencies, an effect visible over a broad range of temporal frequencies. In contrast, at high spatial frequencies, the luminance modulations from blinks are comparable in amplitude to those continually delivered during fixation by ocular drift.

To understand the efficacy of these input signals in driving perceptual responses, we estimated the total power delivered by blinks within the range of temporal sensitivity of the visual system. The distribution in Fig. 1C was here weighted by the temporal function of human contrast sensitivity^49^ and integrated across temporal frequencies. Blinks increase the strength of input signals up to approximately 5 cycles/deg (Fig. 1D). These considerations suggest that, like eye movements^31^, blinks may improve visibility by delivering luminance modulations that effectively increase the contrast of the stimulus on the retina. This effect is expected to occur selectively at low spatial frequencies (Fig. 1E) and to not depend on motor signals associated with the active production of eye blinks.

We tested these predictions in a discrimination task (Fig. 2*A*). Subjects reported whether a 3-cpd grating was rotated by 45° clockwise or counter-clockwise. The spatial frequency of the grating was chosen around the peak of human sensitivity, well within the range where blink transients are predicted to be beneficial. In two separate conditions, subjects were cued to blink either during the presentation of the stimulus (Stimulus-Blink condition) or before its appearance (No-Stimulus-Blink condition; namely, no blink during stimulus presentation). Great care was paid to limit sources of temporal modulations other than eye blinks, both by slowly ramping up the contrast of the stimulus and by discarding all trials in which subjects performed saccades of any amplitude, including microsaccades, during stimulus presentation.

As expected, all subjects were able to blink reliably when prompted, resulting in reaction times and blink characteristics comparable to those previously reported in the literature^9, 10, 43^. The average reaction time ± standard deviation across participants was 389±93 ms (Fig. 2B), and the average blink duration was 165 ± 61 ms (Fig. 2C), which effectively shortened by *∼*15% the exposure to the stimulus at maximum contrast. Blink characteristics were virtually identical in the two conditions of No-Stimulus-Blink and Stimulus-Blink.

As shown in Fig. 2D-E, eye blinks occurring during the examination of the stimulus were beneficial to the task. Despite the reduction in stimulus exposure resulting from the blinks themselves, proportions of correct responses were significantly higher when blinks occurred during the presentation of the stimulus than before its appearance (*p* = 0.010, paired *t*-test). Similar results were also obtained by quantifying performance in terms of the discrimination sensitivity index *d*^t^, with an average improvement of 0.59 (*p* = 0.011, paired *t*-test). This perceptual enhancement was consistent across participants and also reached statistical significance in the individual data from 4 subjects (correct responses: *p <* 0.034; one-tailed *Z*-test corrected for continuity; *d*^t^: *p <* 0.050, bootstrap one-tailed *Z*-test).

**Figure 2:**
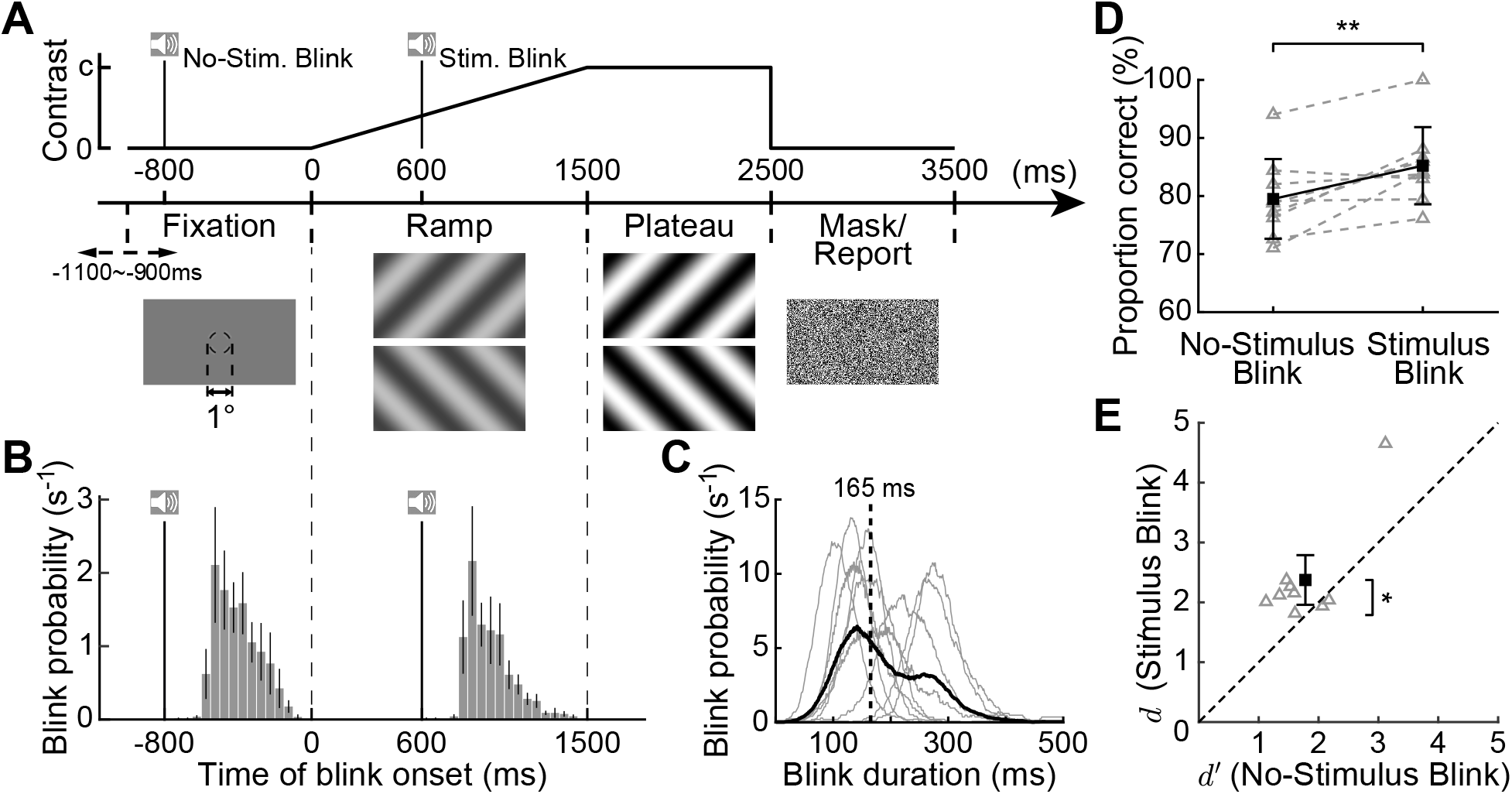
Experimental paradigm and behavioral results. (***A***) Subjects were asked to report the orientation (±45°) of a full-field (21.2° × 11.9°) grating displayed in the presence or absence of blinks. A trial started with the subject fixating at the center of the monitor for a random interval (900-1,100 ms). The stimulus then appeared, ramping up in contrast up to an individually-selected value. A white-noise mask ended the trial. A 50-ms beep instructed subjects to blink either during stimulus presentation (Stimulus-Blink condition; beep 600 ms after stimulus onset) or during the initial period of fixation before the stimulus presentation (No-Stimulus-Blink condition; beep 800 ms before stimulus onset). (***B***, ***C***) Blink characteristics. (***B***) Probability distributions of the timing of blink occurrence in the two conditions. Data represent blink starting times averaged across observers (*N* = 9). The solid vertical lines mark the onset of the auditory cues. Error bars represent ± SEM. (***C***) Probability distributions of blink durations. Both the data from individual subjects (thin gray lines) and the resulting average across observers (bold black line) are shown. The dashed vertical line marks the average blink duration. (***D***, ***E***) Comparison of performance in the presence and absence of blinks during stimulus presentation. The two panels show proportions of correct responses (***D***) and *d*^t^ (***E***). Both averages across subjects (black) and individual subjects data (gray) are shown. Error bars represent ± STD in ***D*** and 95% confidence intervals in ***E*** (** *p* = 0.010, * *p* = 0.011; paired *t*-test; *N* = 9).

These results provide support to the proposal that blink transients are beneficial. To confirm that blinks indeed increased the strength of visual input signals, we estimated the power spectrum of the spatiotemporal luminance flow experienced by the subjects in our experiment. In each trial, we reconstructed the luminance signals resulting from observing the stimulus in the presence of the recorded eye movements and, in the Stimulus Blink trials, the occurrence of an eye blink. In the absence of blinks, the power of the luminance modulations resulting from ocular drifts decreased approximately proportionally to temporal frequency, as previously reported in the literature^48^. In keeping with the predictions of Fig. 1C, eye blinks significantly increased power over a broad band of temporal frequencies (Fig. 3A), effectively resulting in a stronger driving input (Fig. 3B). On average, power increased by approximately 22% across subjects (*p <* 0.00002, paired *t*-test), an effect that also reached significance in the individual data from 6 observers (*p <* 0.042, unpaired one-tailed *t*-test).

**Figure 3:**
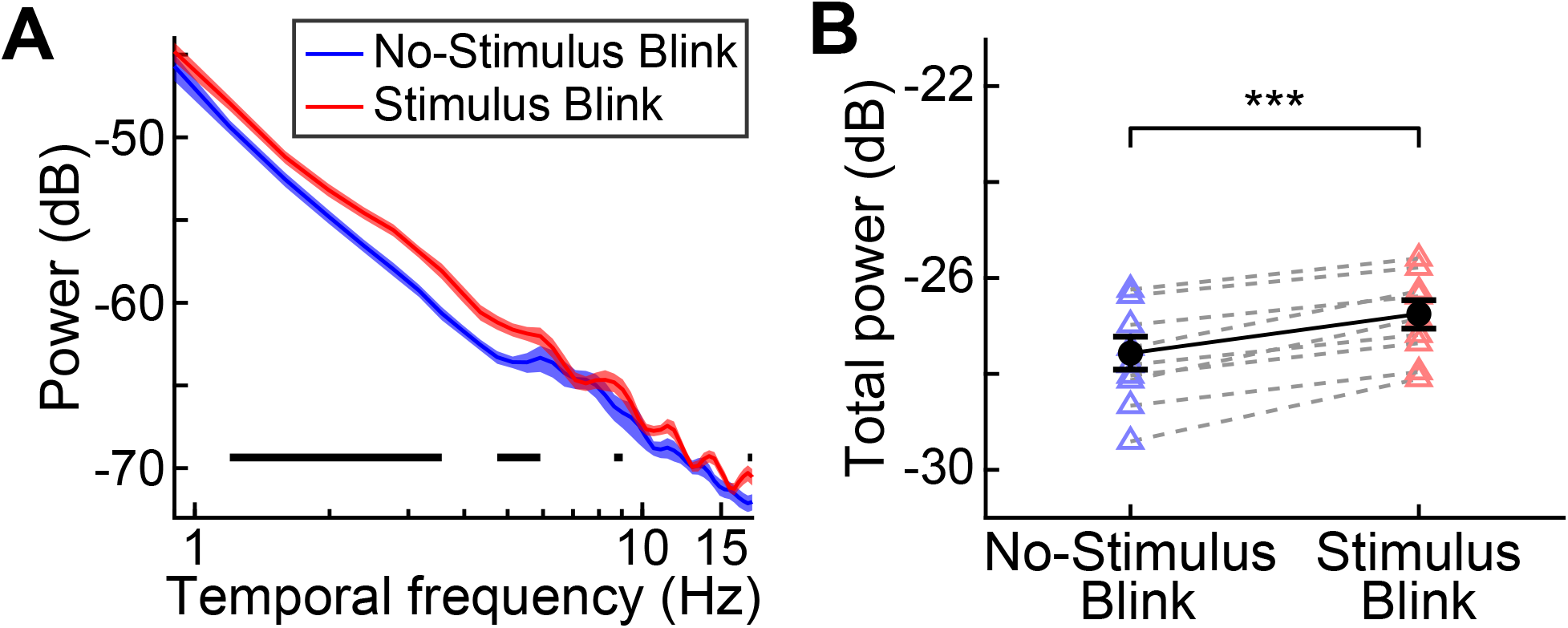
Consequences of eye blinks on visual input signals. (***A***) Power spectra of the spatiotemporal luminance flows experienced by subjects in the two experimental conditions of Fig. 2. Data represent the average spectral density of the visual input resulting from observing the stimulus with the recorded sequences of eye movements and blinks. The occurrence of a blink during stimulus presentation significantly increased power. The horizontal black bars mark statistical significance (*p <* 0.05; paired *t*-test, Bonferroni corrected). Shaded regions represent *±* SEM. (***B***) Strength of input signals within the temporal range of human sensitivity. The power in ***A*** is here weighted by the average human contrast sensitivity function and integrated across temporal frequency. Circles and triangles show averages across subjects and individual subjects’ data, respectively. Error bars represent *±* SEM (*** *p <* 0.00002, paired *t*-test).

The model in Fig. 1 makes a more specific prediction: if the transients delivered by eye blinks are indeed responsible for the perceptual enhancement shown in Fig. 2, one may expect this improvement to be confined to low spatial frequencies, the range in which the modulations from fixational eye movements are small. To test this hypothesis, we repeated the experiment using gratings at either low (1 cpd) or high (10 cpd) spatial frequency. Apart from the stimulus change, the task and procedures were otherwise identical to those described in Fig. 2. As before, with a low spatial frequency stimulus, the occurrence of an eye blink enhanced performance (Fig. 4A). In contrast, blinks had no effect during viewing of the high-frequency grating, and discrimination performance was virtually identical when blinks occurred before and during stimulus presentation. In keeping with these findings and the predictions of Fig. 1, blinks also increased the efficacy of the visual signals impinging onto the retina during exposure to a 1-cpd grating (*p ≤* 0.0006, unpaired *t*-test) but not with a 10-cpd grating (*p ≥* 0.141, unpaired *t*-test; Fig. 4B). Thus, these results corroborate the idea that the perceptual consequences of blinks measured in our experiments originated from their luminance transients.

**Figure 4:**
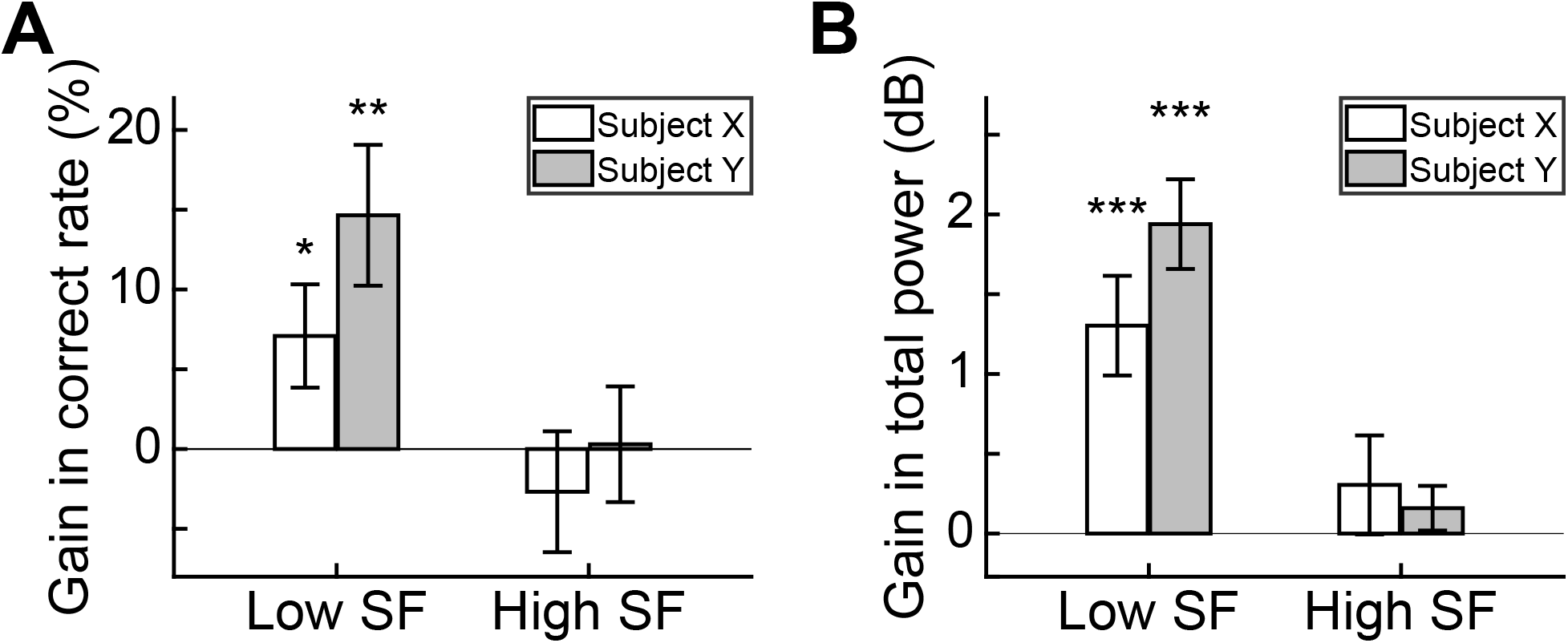
Eye blinks selectively benefit low spatial frequencies. (***A***) Changes in performance between Stimulus-Blink and No-Stimulus-Blink conditions during the presentation of a 1-cpd (low spatial frequency) or 10-cpd grating (high spatial frequency). Data from two subjects are shown in separate bars. Error bars represent ± STD. * *p* = 0.042, ** *p* = 0.002; *Z*-test corrected for continuity. (***B***) Gain in the power of retinal stimulation resulting from blinking. Data represent the ratios of power delivered within the temporal range of human sensitivity by the eye movements and blinks recorded in the two experimental conditions. *** *p <*= 0.0006; unpaired *t*-test. Error bars indicate ± SEM.

Our predictions are purely based on the characteristics of the visual signals delivered by blinks: these abrupt transients redistribute the spatial power of the stimulus across temporal frequencies, effectively yielding a stronger driving input at low spatial frequencies. These considerations imply that the visual system should benefit from these transients irrespective of their origin, *i.e.*, whether caused by blinks or generated from the external stimulus. To investigate this hypothesis, in a third experiment, rather than instructing subjects to actively blink, we passively exposed them to reconstructions of blink transients obtained by directly modulating the luminance of the stimulus on the display (Fig. 5A).

**Figure 5:**
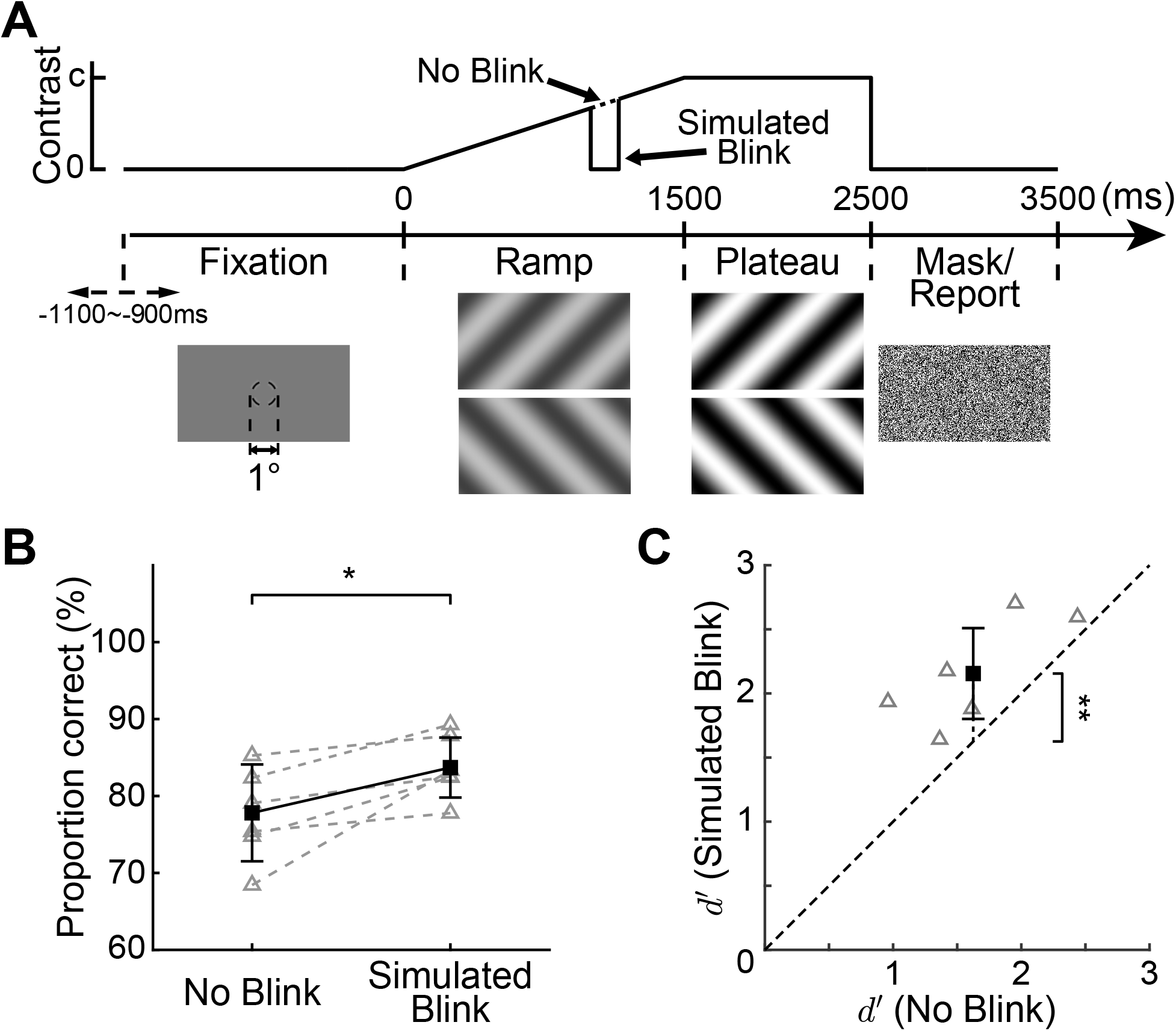
Perceptual consequences of simulated blinks. (***A***) In an experiment similar to that of Fig. 2A, subjects were exposed to changes in the stimulus that resembled the luminance transients elicited by eye blinks. Each simulated blink consisted of a brief period of stimulus blanking with onset and duration randomly sampled from the previously collected individual blink data. (***B***, ***C***) Comparison of performance in the presence and absence of a simulated blink. The two panels show proportions of correct responses (***B***) and *d*^t^ (***C***). Both averages across subjects (*N* = 6; black) and individual subjects’ data (gray) are shown. Error bars represent ± STD in ***B*** and 95% confidence intervals in ***C*** (\*\**p* = 0.012, \**p* = 0.023; paired *t*-test).

The results in Fig. 5 show that exposure to simulated blinks is beneficial. Performance was significantly higher in the simulated-blink trials than in the absence of the abrupt luminance changes, an effect evident both in the percentages of correct responses (*p* = 0.023, paired *t*-test; Fig. 5B) and in the discriminability index (average improvement of 0.53; *p* = 0.012, paired *t*-test; Fig. 5C). The extent of the improvement obtained with simulated blinks was comparable in magnitude to that measured with real eye blinks (*p* = 0.820 and *p* = 0.803 for correct rate and *d*^t^, respectively; unpaired *t*-test). This enhancement was highly consistent across subjects: all participants exhibited higher performance in the simulated-blink condition, as directly visible in their individual data (Fig. 5B,C). Thus, passive exposure to brief luminance transients similar to those resulting from eye blinks enhances visual sensitivity.

## Discussion

Humans blink their eyes frequently during normal viewing, more often than it seems necessary for refreshing the tear film^6, 7, 19^. Since each blink lasts 100-300 ms, it is estimated that an individual can spend as much as 10% of their awake time while blinking^52^. It has long been questioned how the visual system deals with the associated interruptions in the input to the retina^8, 9, 53^ and whether eye blinks serve other functions besides lubricating the eye^4, 5^. The results of this study provide a possible answer to these questions: rather than impairing visual processing as commonly assumed, blinks enhance sensitivity via their luminance transients.

Specifically, our data show that (a) discrimination of spatial patterns is facilitated when blinks occur during stimulus presentation (Fig. 2); and (b) blink transients deliver powerful luminance modulations within the temporal range of visual sensitivity, increasing the strength of input signals relative to sustained fixation (Fig. 3). These effects only occur for stimuli at sufficiently low spatial frequencies (Fig. 4), as predicted from a model of the spatiotemporal luminance flow to the retina (Fig. 1). Furthermore, they occur irrespective of whether the abrupt luminance transients are actively generated by blinks or passively experienced (Fig. 5), an observation that controls for possible extraretinal contributions as well as changes in retinal image quality with blinks. The resulting perceptual improvements are considerable and sufficient to overcome the temporal loss in stimulus exposure in our experiments.

At first sight, our findings appear to conflict with the considerable evidence of a perceptual suppression accompanying blinks^8, 9, 11^. Ingenious experiments that projected light directly onto the retina through the mouth^8^ or used specula to keep the eyelids open^11, 54^ have revealed an attenuation in sensitivity that precedes and outlasts the blink itself^13^. These studies, however, focused on a different question than the one investigated here: how blinks affect the visibility of transient events, such as a briefly displayed probe. These events are rare under natural viewing conditions, and their luminance modulations interact with those normally resulting from blinks. In contrast, here we focused on how blinks influence the representation of a stationary scene continuously present throughout the blink, as it normally occurs outside the laboratory. To this end, in our experiments, we were careful to minimize all transients other than those caused by eye movements and the blinks themselves.

In fact, our results provide a possible explanation for the reports of previous studies that did not temporally manipulate stimuli, including the observations that blinks counteract and prevent image fading during prolonged fixation^55, 56^. This effect is what one would expect from a visual system that relies on blinks transients to enhance sensitivity to low spatial frequencies. Our results are also consistent with the neural modulations measured at the time of eye blinks. Intracranial electrocorticographic recordings in humans^26^ indicate that the changes in visual stimulation elicited by blinks sharply modulate neural responses, with an initial drop in activity followed by a strong overshoot at stimulus reappearance. This signal is consistent with the spatiotemporal redistribution of power in Fig. 3, which predicts strong responses when the eyes reopen.

It is worth pointing out that the performance improvement measured in our study differs from the attentional influences previously reported in the literature^21^. Blinks have been found to be beneficial in rapid serial visual presentations (RSVP), in which subjects identify target stimuli in random streams of distractors. This effect has been attributed to an attentional facilitation following blinks^20^ and, unlike the enhancement we observed, only occurs with real blinks, not simulated ones. In fact, this facilitation cannot be caused by the luminance modulations delivered by blinks, which, in an RSVP procedure, are swamped by the transients resulting from the rapid succession of images. In contrast, the perceptual enhancement measured in our experiments follows the structure of blink-induced luminance modulations: it occurs with both real and simulated blinks; and is present at low but not high spatial frequencies.

The finding that the visual system takes advantage of blinks’ modulations acquires further importance in the context of visual perception theories arguing for a temporal encoding of spatial information^27, 57–61^. During natural viewing, the retina is continually exposed to changes in luminance, as saccades alternate with fixational eye movements^38–40^. This behavior appears to confound the processing of spatial information^14, 15, 18, 62, 63^. However, evidence is increasingly accumulating for an alternative viewpoint, one that casts a more positive light on these input changes^31^. According to this view, spatial information is not just conveyed by the location of neurons within maps, as commonly assumed, but also by the way motor behavior shapes the temporal structure of neural responses^27–29^. This idea builds on the observation that the resulting visual input signals stimulate the retina with temporal changes rich in spatial information, which critically depend on how the eye moves.

A number of findings from our laboratory support this proposal of active space-time encoding. We showed not only that fixational eye movements critically enhance—rather than degrade—vision of fine spatial detail^30, 36, 44^, but also that their luminance modulations are matched to the statistics of natural scenes^48^, forming a continuum with the modulations from larger eye movements^41^. Furthermore, we discovered a previously unexpected level of control in fixational eye movements^36, 64–66^, which appears essential for reaching high visual acuity and is consistent with its consequence on the retinal stimulus. The current study adds to this previous body of work by showing that the strong luminance transients exerted by eye blinks are also useful. Like saccades, the luminance signals from blinks enhance low spatial frequencies relative to ocular drift and possess temporal frequencies that are expected to strongly activate retinal neurons^67^. However, unlike eye movements, blinks modulate luminance equally at all spatial frequencies, thus not counterbalancing the power spectrum of natural scenes. This implies that blinks deliver much stronger signals than saccades at low spatial frequencies.

In the context of theories advocating for a temporal encoding of spatial information, the notion of blink suppression acquires an alternative conceptual interpretation. Like the attenuation in sensitivity that accompanies saccades^13, 15, 18^, blink suppression is believed to serve the purpose of reducing visibility to abrupt changes in retinal stimulation, thus facilitating the establishment of stable visual representations. However, the possibility emerges that, rather than mechanisms for suppressing sensitivity to sensory changes, these phenomena may actually be corollaries of using brief probes to test neural pathways tuned to extract information from luminance transients. That is, these effects may reflect the interference between the neural responses to the probes used to measure sensitivity and the process of extracting spatial information from oculomotor-induced input transients. This hypothesis is consistent with the observations that both saccade and blink suppression are stronger for low spatial frequency^11^ and achromatic stimuli^54^, which is the type of information expected to be conveyed by saccade and blink transients.

In sum, we have shown that the luminance transients resulting from eye blinks enhance contrast sensitivity to low spatial frequencies, an effect that occurs despite the loss of exposure imposed by the blink itself. Thus, in addition to lubricating the eye, blinks also appear to serve an information-processing function by shaping the spatial content of the signals that fall within the temporal range of visual sensitivity. It is known that human plan blinks to avoid missing salient events^68^. Further work is needed to investigate whether blinks are also strategically executed according to the role played by their transients in the acquisition of visual information.

## Acknowledgement

This work was supported by NIH grants R01 EY18363 and P30 EY001319.

